# Identification of novel *cis*-acting elements, E-box and ISRE, regulating IFNγ-IRF1 axis-mediated NLRC5 expression

**DOI:** 10.1101/2025.02.20.639218

**Authors:** Ji-Seung Yoo, Yusuke Kasuga, Saptha Vijayan, Suhail Yousuf, Koichi S. Kobayashi

## Abstract

NLRC5 and CIITA are the primary transcriptional regulators of MHC class I and MHC class II, respectively, and play essential roles in adaptive immunity. While the regulatory mechanisms of CIITA have been extensively characterized, the transcriptional control of NLRC5 remains incompletely understood. In this study, we identified two novel conserved *cis*-regulatory elements within the NLRC5 promoter, an E-box and an ISRE. Furthermore, we revealed IRF1 as a novel transcriptional regulator of NLRC5, binding directly to the ISRE within the NLRC5 promoter. Using the newly identified NLRC5 ISRE, we established a screening platform to identify modulators of the IFNγ-IRF1-NLRC5 axis. This system corroborated the inhibitory effects of known viral antagonists and led to the identification of novel SARS-CoV-2 viral factors that suppress IFNγ- mediated IRF1 nuclear translocation, thereby inhibiting the MHC class I pathway. By elucidating the previously unproven molecular mechanism underlying IFNγ-mediated NLRC5 regulation, our study provides critical insights into viral immune evasion strategies and the modulation of antigen presentation. These findings may facilitate the development of MHC class I-targeted therapeutics by modulating the IRF1-NLRC5 axis.

## INTRODUCTION

The major histocompatibility complex (MHC) class I antigen presentation pathway is essential for host defense against intracellular pathogens and cancer. By presenting immunogenic antigens to CD8+ cytotoxic T cells, the MHC class I program facilitates the targeted elimination of pathogen-infected cells or cancer cells. Type I and II interferons (IFNs), are critical in activating the MHC class I pathway. IFN stimulation induces signal transducer and activator of transcription 1 (STAT1) activation, which subsequently upregulates interferon regulatory factor 1 (IRF1) and NOD-like receptor family CARD domain containing 5 (NLRC5), cooperatively enhancing the expression of MHC class I pathway components^1^.

NLRC5, a member of the NOD-like receptor family, functions as the MHC class I transactivator (CITA) and serves as a master transcriptional regulator of genes within the MHC class I pathway. Under conditions of infection or cancer, NLRC5 undergoes conformational changes facilitated by ATP binding to its NACHT domain, enabling its nuclear translocation via a bipartite nuclear localization signal (NLS). Then NLRC5 interacts with multiple transcription factors at the MHC class I promoter, forming the CITA enhanceosome, a functional protein-DNA complex critical for the activation of MHC class I gene expression^2^.

As a central regulator of the MHC class I antigen presentation pathway, NLRC5 is a primary target for immune evasion mechanisms in both cancer and infectious diseases. Tumor cells frequently downregulate NLRC5 expression through genetic and epigenetic alterations, including promoter methylation, mutations, and copy number loss. These modifications impair MHC class I expression, compromise CD8+ T-cell recruitment, and correlate with adverse clinical outcomes^3^. Intracellular pathogens have also evolved strategies to suppress NLRC5-mediated immune responses. Our previous study showed that severe acute respiratory syndrome coronavirus 2 (SARS-CoV-2) encodes open reading frame 6 (ORF6), which suppresses NLRC5 expression by interfering with IFN-STAT-IRF1 signaling, leading to reduced MHC class I gene transcription^4^. Furthermore, ORF6 prevents NLRC5 nuclear translocation, thereby inhibiting its function as a CITA^4^. Similarly, recent study showed that *Orientia tsutsugamushi* utilizes the ankyrin repeat (AR)- containing effector protein 5 (Ank5) protein to obstruct NLRC5 nuclear translocation and promote its degradation^5^. Given the critical role of NLRC5 in immune defense, the precise regulation of its expression is essential. However, the molecular mechanism underlying its transcription remain poorly understood.

In this study, we identified two novel *cis*-acting elements within the NLRC5 promoter, the enhancer box (E-box) and the interferon-sensitive response element (ISRE). We clarified that IRF1 acts as a specific transcription factor for NLRC5, directly binding to the ISRE and regulating NLRC5 gene expression, thereby facilitating MHC class I pathway activation.

Furthermore, we developed a novel screening system by integrating the NLRC5 ISRE with reporter genes, enabling the identification of antagonists targeting the IRF1-NLRC5 axis. Using this approach, we discovered a previously unknown immune evasion strategy employed by SARS-CoV-2. Additionally, this system provides a powerful tool for screening potential inhibitors that target the IRF1-NLRC5 axis-mediated MHC class I expression. Altogether, our findings provide new insights into NLRC5 transcriptional regulation and suggest potential directions for the development of targeted therapeutic approaches.

## RESULTS

### IFN-dependent and -independent regulation of NLRC5 expression

Previous studies have demonstrated that *NLRC5* gene expression is induced by both Type I and Type II interferons (IFNs)^6^. To further explore the molecular mechanisms underlying the transcriptional regulation of NLRC5, we developed a luciferase reporter assay system based on a previously characterized NLRC5 promoter region^7^, with an additional downstream extension of 147 nucleotides to include the Exon 1 domain (Figure S1A). First, we assessed the activity of this reporter construct for its responsiveness to type I and type II IFNs. As expected, both types of IFNs significantly enhanced NLRC5 promoter activity (Figures S1B and S1C). Interestingly, even in the absence of IFN stimulation, promoter activity was significantly increased by approximately 5-6-fold compared to the empty control (pGL3), suggesting the involvement of an IFN-independent mechanism in NLRC5 promoter activation. This observation was further corroborated using IFNAR1 KO HeLa cells stimulated with PAMP RNAs, including polyinosinic-polycytidylic acid (pI:C) and 5’-triphosphate RNA (5’ppp) (Figure S1D). Collectively, these results indicate that *NLRC5* gene expression is regulated through both IFN-dependent and IFN-independent mechanisms.

### IRF1 is a transcriptional regulator of *NLRC5* gene

Antiviral IFN systems coordinate secondary innate immune responses by establishing an antiviral state through the JAK-STAT pathway. These responses are triggered by primary immune signaling, which is activated by innate immune sensors and transcription factors^8^. Among these, IRF1, IRF3, and IRF7 are crucial transcriptional regulators that drive the activation of antiviral IFN signaling pathways^9^.

To elucidate the mechanism regulating NLRC5 expression before IFN production, we investigated the potential contribution of key antiviral signaling-related transcription factors, including IRF1, IRF3, and IRF7, to NLRC5 gene expression. Using both WT and IFNAR1 KO HeLa cells, we assessed NLRC5 promoter activity following the overexpression of these transcription factors. Surprisingly, IRF1 overexpression markedly increased NLRC5 promoter activity in both WT (Figure 1A) and IFN-deficient (Figure 1B) systems, highlighting its pivotal role in regulating NLRC5 expression independently of IFN production.

**Figure 1.**
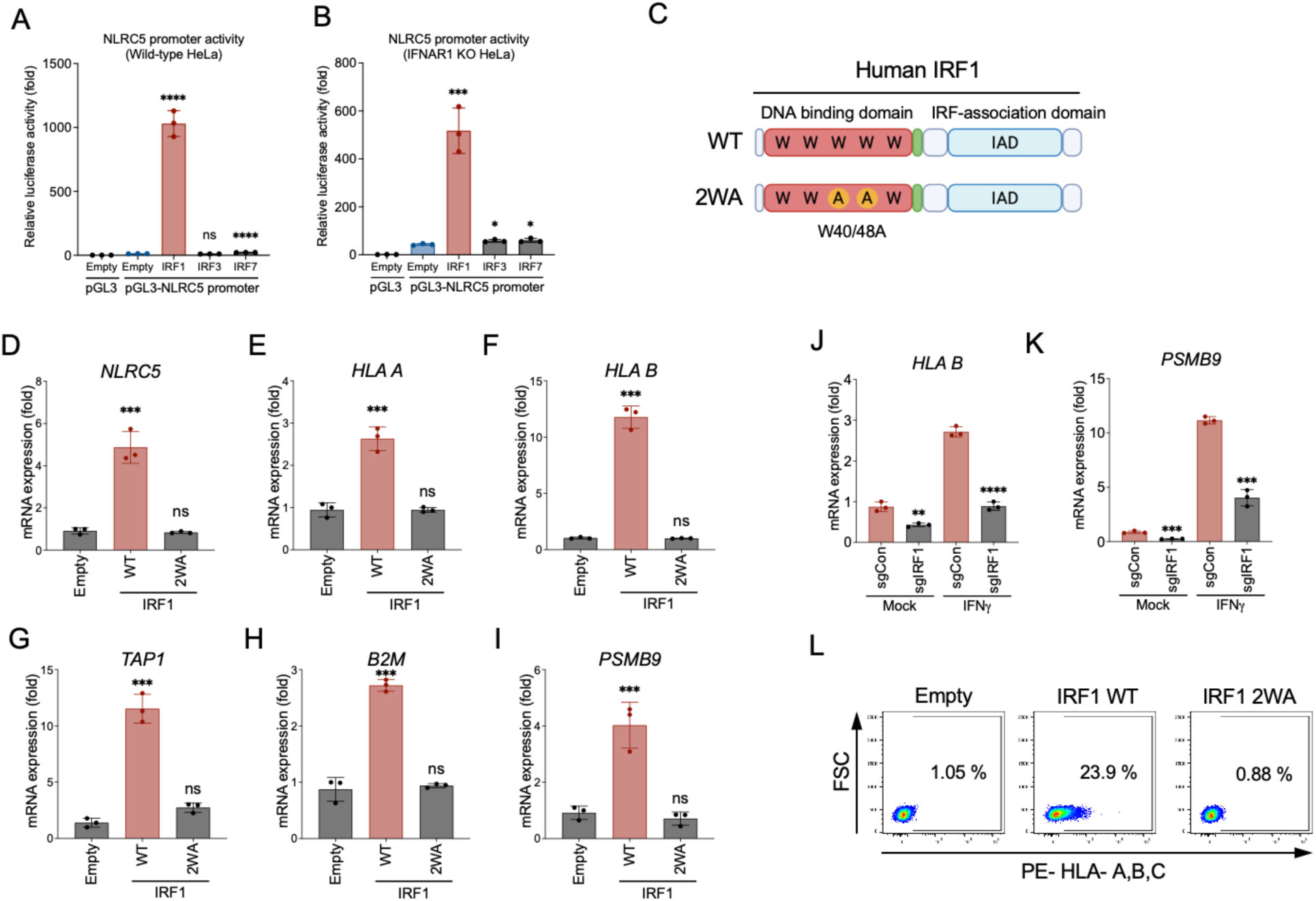
IRF1 is a transcriptional regulator of NLRC5. (A and B) NLRC5 promoter activity was assessed using a luciferase-based reporter assay in HeLa WT (A) and IFNAR1 KO (B) cells, induced with empty control, IRF1, IRF3, and IRF7 to monitor NLRC5 promoter activity. (C) Schematic depiction of IRF1 WT and 2WA constructs. (D to I) RT-qPCR analysis was conducted to evaluate the expression of MHC class I-related genes in HeLa cells. The expression levels of *NLRC5* (D), *HLA A* (E), *HLA B* (F), *TAP1* (G), *B2M* (H), and *PSMB9* (I) genes mediated by WT IRF1 were presented as fold changes relative to the empty control or IRF1 2WA mutant. (J and K) IFNγ-induced expression of *HLA B* (J) and *PSMB9* (K) genes was analyzed in either control (sgCon) or IRF1 KO (sgIRF1) PH5CH8 cells using RT-qPCR. (L) Surface expression levels of HLA-A, B, and C were assessed by flow cytometry in HEK293T cells expressing either empty vector, IRF1 WT, or IRF1 2WA constructs.

Next, we investigated the IRF1-induced transcriptional regulation of *NLRC5* and genes associated with the MHC class I pathway. To serve as a negative control, we generated a functionally inactive IRF1 mutant by substituting two tryptophan residues in its DNA-binding motif with alanines (2WA) (Figure 1C). The gene expression of *NLRC5* and other genes in the MHC class I pathway, including human leukocyte antigen A (*HLA-A*), *HLA-B*, transporter associated with antigen processing 1 (*TAP1*), beta-2 microglobulin (*B2M*), and proteasome subunit beta type-9 (*PSMB9*), was significantly induced by the overexpression of WT IRF1, but not by the 2WA mutant (Figures 1D-1I). Furthermore, IRF1 KO cells displayed defects in IFNγ-induced *HLA-B* and *PSMB9* gene induction (Figures 1J and 1K). Notably, since IRF1 did not affect the expression of other known transcriptional regulators of the MHC class I pathway (Figure S2), its role in the MHC class I pathway appears to be highly specific to NLRC5. While overexpression of WT IRF1 strongly enhanced the surface expression of HLA proteins, the IRF1 2WA mutant failed to do so (Figure 1L), further corroborating the critical role of IRF1 in the transcriptional regulation of the NLRC5- MHC class I axis.

### Identification of the *cis*-regulatory element in the NLRC5 promoter

Previously, Kuenzel and colleagues reported two putative STAT binding sites within the upstream region of the NLRC5 gene, κB/STAT (-1365 to -1328) and STAT (-477 to -442)^7^. We revisited their work to explore the IRF1-interacting region in the NLRC5 promoter. We constructed a series of luciferase-based reporter genes in which one or both reported STAT binding elements were deleted. Additionally, we generated reporter constructs that included or excluded the partial Exon 1-containing region (-73 to +39) (Figure 2A).

**Figure 2.**
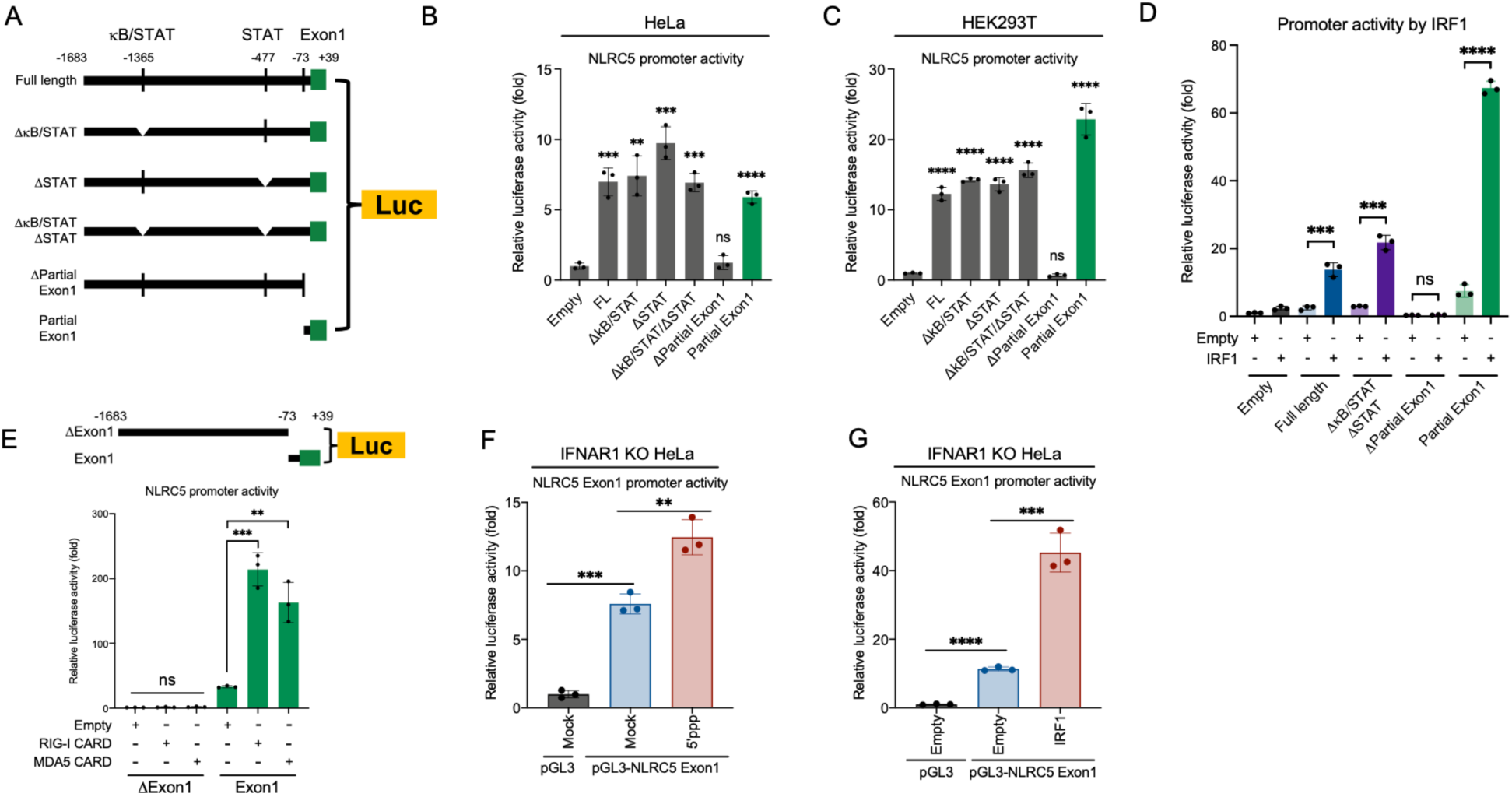
Identification of the *cis*-regulatory element for NLRC5 expression (A) Schematic illustration of the luciferase-based NLRC5 promoter reporter constructs. (B and C) NLRC5 promoter activity was assessed by luciferase assay in HeLa (B) and HEK293T (C) cells. (D) NLRC5 promoter activity was measured by luciferase assay in HEK293T cells overexpressing IRF1 or treated with an empty vector. (E) NLRC5 promoter activity was monitored in HEK293T cells induced with either empty, RIG-I-CARD, or MDA5-CARD constructs. Schematic illustration of reporter constructs used for this analysis is shown above. (F and G) NLRC5 promoter activity was evaluated in IFNAR1 KO HeLa cells stimulated with 5’-ppp RNA (F) or through IRF1 overexpression (G).

In contrast to previous findings, the deletion of two STAT binding sites did not impact NLRC5 promoter activity in HeLa (Figure 2B) and HEK293T (Figure 2C) cells. Interestingly, however, the removal of a partial Exon 1 region completely abolished luciferase activity. Conversely, a construct containing only the partial Exon 1 region was sufficient to drive NLRC5 promoter activity. The observed activity levels were comparable to, or even exceeded, those of the full-length control (Figures 2B and 2C). These results suggest that a *cis*-regulatory element critical for NLRC5 promoter activity is located near the Exon 1 region.

Next, we examined the impact of IRF1 on the promoter activity of these reporter constructs. Overexpression of IRF1 significantly increased the promoter activity of both the full-length construct and the construct with double-deleted STAT binding sites. In contrast, the construct lacking the partial Exon 1 region exhibited a complete loss of luciferase activity. Interestingly, the construct containing only the partial Exon 1 region showed a dramatic induction of IRF1-mediated luciferase activity (Figure 2D). These results strongly suggest that the region adjacent to Exon 1 functions as a critical *cis*-regulatory element for IRF1 activity.

Constitutively active forms of two cytoplasmic RNA virus sensors, RIG-I CARD-only and MDA5 CARD-only, which mimic viral infection conditions^10,11^, significantly enhanced the promoter activity of the construct containing only the partial Exon 1 region (Figure 2E). This finding indicates that certain downstream factor of RIG-I-like receptor (RLR) signaling may contribute to the transcriptional regulation of the NLRC5 gene. This hypothesis was further validated in IFNAR1 KO HeLa cells stimulated with either 5’-ppp RNA (Figure 2F) or IRF1 overexpression (Figure 2G), confirming again that IRF1 (Figures 1A and 1B), rather than interferon system, acts as the key downstream factor of RLR signaling for the early induction of the NLRC5 gene.

### Identification of the E-box and ISRE in the NLRC5 promoter

Since our data demonstrate that IRF1 enhances NLRC5 promoter activity, we sought to identify potential ISRE that serve as recognition sites for IRF1. Using BLAST analysis, we examined the NLRC5 promoter sequences (-73∼+39; based on human DNA sequence) (Figure 2A) across mammals to identify conserved regions. Interestingly, two domains within the NLRC5 promoter region were found to be perfectly conserved across all mammals analyzed (Figure 4A). The sequence ’CACGTG’ is a well-known E-box motif (CANNTG, where N represents any nucleotide) that plays a pivotal role in gene expression^12^ while ’CTTTCAGTTTC’ corresponds to a typical ISRE, where IRF1 binds to regulate gene activity^13^. Notably, both the E-box and ISRE are located within the promoter region of the class II major histocompatibility complex (MHC class II) transactivator (CIITA) and play a key role in regulating CIITA gene expression^14,15^. Importantly, the DNA sequences near NLRC5 Exon 1 are predicted to reside within a CpG island (Figure S3A). Moreover, experimentally validated transcription start site (TSS) data confirmed that NLRC5 gene transcription initiates from the ISRE site we identified (Figures S3B and S3C). Based on this result, we term the juxtaposed enhancer that activates the ISRE within the Exon 1 region the ’<u>J</u>uxtaposed enhancer-box and ISRE (JI; -73 to +6 position)’ domain.

Among the DNA sequences analyzed in the BLAST search, we selected the mouse DNA sequence corresponding to the human JI domain and generated a reporter construct. Both human and mouse JI domain constructs showed similar increases in promoter activity upon IRF1 overexpression, providing evidence that these regions harbor essential *cis*-acting elements required for NLRC5 gene expression (Figure 3B).

**Figure 3.**
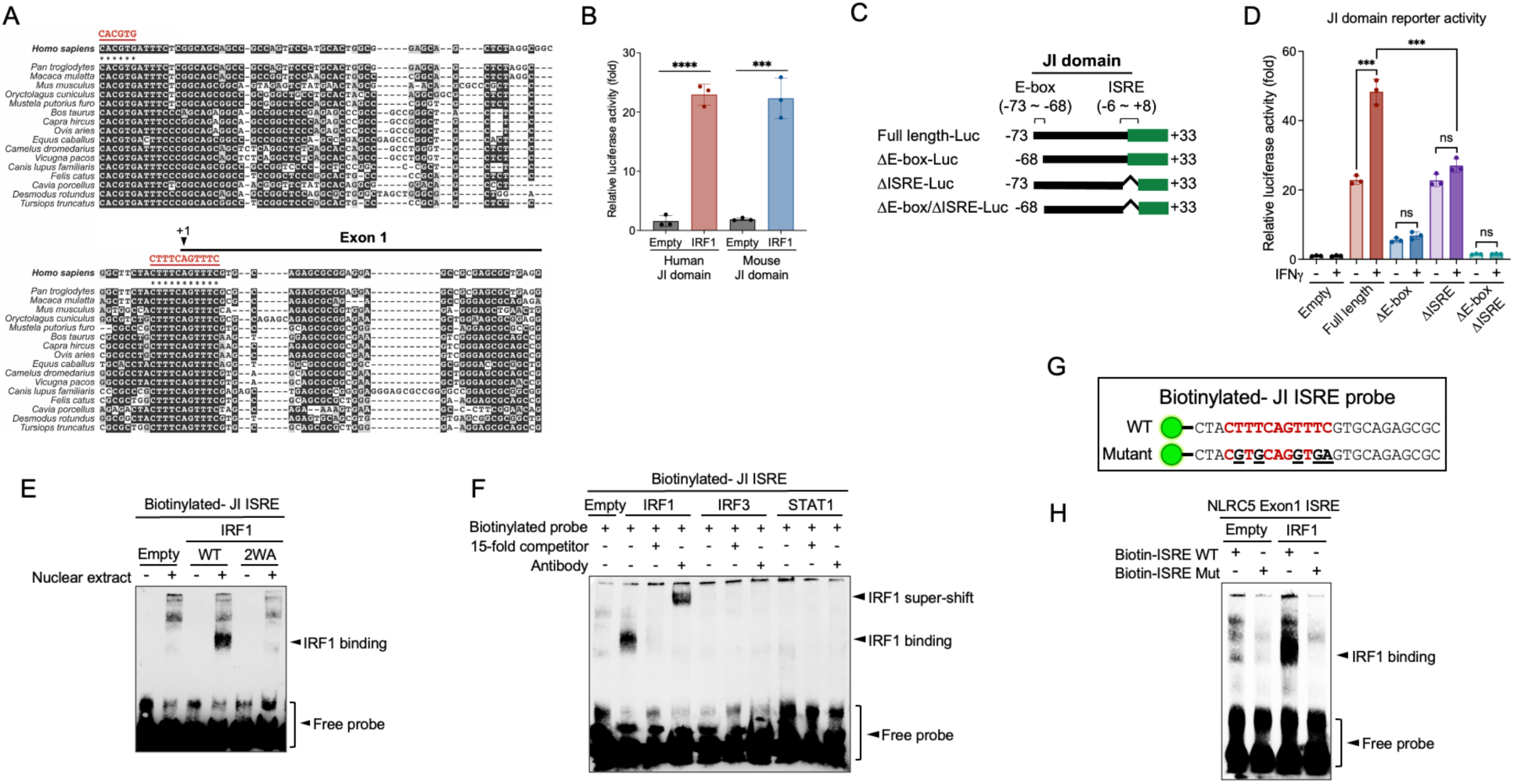
Identification of E-box and ISRE in the NLRC5 promoter (A) DNA sequence alignment of the NLRC5 promoter regions across various vertebrates, with E-box and ISRE regions highlighted in red. (B) The promoter activity of a NLRC5 JI domain was assessed by luciferase assay in HEK293T cells transfected with either an empty vector or IRF1 expression construct. (C) Schematic illustration of the luciferase-based NLRC5 JI domain reporter constructs, with the exact nucleotide positions indicated by numbers. (D) NLRC5 JI domain promoter activity was measured by luciferase assay in HEK293T cells, with or without IFNγ treatment. (E to H) EMSA analysis using biotinylated probes. (E) Complex formation between the JI ISRE probe and IRF1 WT or 2WA was analyzed by EMSA. (F) The complex formation of the JI ISRE probe with IRF1, IRF3, and STAT1 was evaluated. Super-shift assays were also performed using anti-IRF1, anti-IRF3, or anti-STAT1 antibodies. (G) Biotinylated JI ISRE WT or mutant probes are shown. (H) Complex formation between the JI ISRE WT or mutant probe and IRF1 was evaluated.

**Figure 4.**
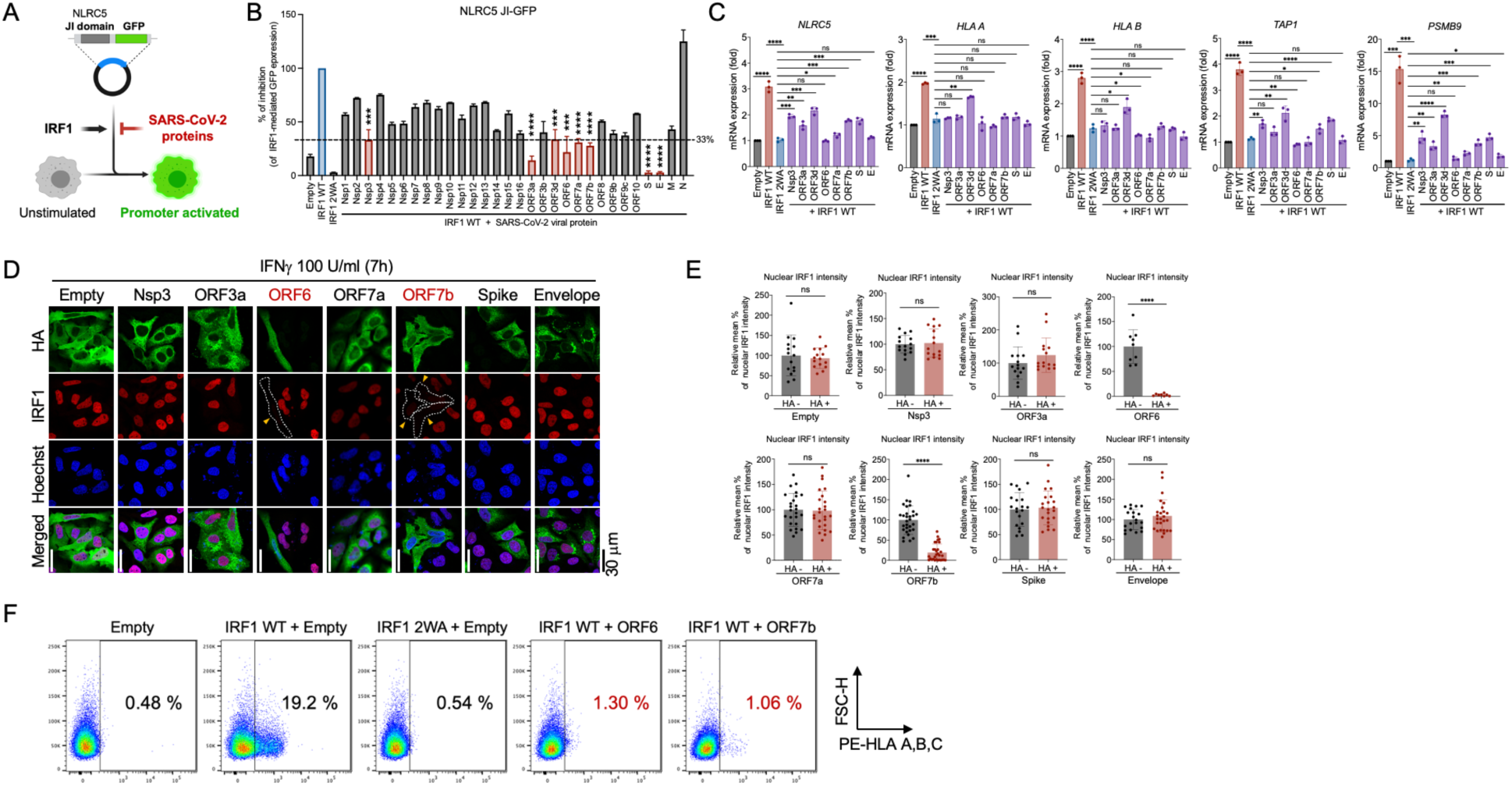
Screening of the SARS-CoV-2 proteins antagonizing the IRF1-NLRC5 axis- derived MHC class I pathway (A) Schematic illustration of the strategy for screening SARS-CoV-2 antagonists using the NLRC5 JI-GFP system. (B) Inhibitory effect of SARS-CoV-2 viral proteins on IRF1-mediated NLRC5 JI- GFP reporter activity, measured by flow cytometry. As controls, empty vector or IRF1 2WA were used. The dotted line indicates a 33% reduction compared to the positive control (IRF1 WT, blue bar). (C) The inhibitory effect of SARS-CoV-2 factors on IRF1-mediated expression of MHC class I-related genes (NLRC5, HLA A, HLA B, TAP1, and PSMB9) was evaluated by RT-qPCR. (D and E) The inhibitory effect of SARS-CoV-2 factors on IFNγ-induced IRF1 nuclear translocation was analyzed by confocal microscopy (D), with quantitative analysis of the image data presented as a bar graph (E). (F) The inhibitory effect of SARS-CoV-2 ORF6 and ORF7b on IRF1-induced HLA A, B, C surface expression was evaluated by flow cytometry.

To further investigate whether the E-box and ISRE within the JI domain are essential for NLRC5 promoter activity, we generated a series of luciferase-based reporter constructs in which the E-box, ISRE, or both elements were deleted (Figure 3C). The full-length JI construct exhibited more than a 20-fold increase in promoter activity compared to the empty control, with further enhancement observed upon IRF1 overexpression (Figure 3D). Interestingly, deletion of the E-box from the JI domain resulted in a dramatic reduction in promoter activity, while deletion of the ISRE led to a similar basal level of promoter activity as the empty control, although it no longer responded to IRF1 stimulation. Furthermore, deletion of both elements from the JI domain completely abolished promoter activity (Figure 3D). These data clearly suggest that the E-box region is critical for maintaining basal NLRC5 promoter activity, while the ISRE is essential for IRF1-dependent regulation of NLRC5 gene expression.

### IRF1 directly binds to the ISRE in the JI domain

To investigate whether IRF1 directly binds to the ISRE in the JI domain, we performed an electrophoretic mobility shift assay (EMSA). A biotinylated probe, consisting of 25 nucleotides from the JI domain containing the ISRE, was synthesized and used in the assay (see **Methods**). Interestingly, a shifted band was observed when the ISRE probe was incubated with WT IRF1- expressed cell lysate. However, no shifted band was detected when incubated with either the empty control or the IRF1 2WA mutant (Figure 3E). Moreover, while the ISRE probe formed a complex with IRF1 and an anti-IRF1 antibody (IRF1 super-shift), incubation with IRF3- and STAT- expressed cell lysates did not yield any shifted bands, confirming that IRF1 specifically interacts with the ISRE in the JI domain (Figure 3F). Next, to determine whether this interaction is indeed sequence-specific, we created a mutant ISRE probe by randomly substituting four thymidines with guanines and one cytidine with an adenosine (Figure 3G). While the WT ISRE probe formed a complex with IRF1, the mutant ISRE probe did not, providing strong evidence that the IRF1-JI ISRE interaction is sequence-specific (Figure 3H).

### Establishing a GFP-based reporter system for monitoring NLRC5 expression

Since NLRC5 is essential for cellular defense mechanisms against infections and tumorigenesis by regulating the MHC class I pathway, we aimed to establish a monitoring system for NLRC5 expression. To this end, we engineered a construct by fusing green fluorescent protein (GFP) or luciferase genes to the downstream region of the JI domain, referred to as JI-GFP or JI-Luc. The expression of GFP was evaluated by overexpressing either IRF1 WT or IRF1 2WA in combination with JI-GFP (Figure S4A). Consistent with the data shown in Figure 3, robust JI-GFP expression was observed in cells overexpressing WT IRF1, while only minimal expression was detected in cells transfected with either the empty vector or 2WA IRF1, as verified by fluorescence microscopy (Figure S4B) and flow cytometry analysis (Figure S4C). Moreover, confocal microscopy demonstrated that activation of the JI-GFP reporter by IRF1 occurs independently of the IFN signaling pathway (Figures S4D and S4E), further supporting our earlier observations in Figures S1 and 2.

### Feasibility of the JI-GFP system in screening MHC class I-targeting viral antagonists

Next, we aimed to determine whether the NLRC5 JI domain-containing reporters (JI-GFP and JI- Luc) are suitable tools for screening MHC class I-targeting viral antagonists. To this end, we tested a panel of well-characterized viral antagonists, including severe acute respiratory syndrome coronavirus 2 (SARS-CoV-2) open reading frame 6 (ORF6)^4^, hepatitis C virus (HCV) core protein^16^, Kaposi’s sarcoma-associated herpesvirus (KSHV) K11^17^, and hepatitis B virus (HBV) X protein^18^. These antagonists were evaluated for their ability to suppress IRF1-driven activation of the JI-Luc reporter (Figure S4F). All tested viral antagonists significantly inhibited JI-Luc reporter activity, corroborating previously reported phenotypes (Figure S4G). These findings were further validated by analyzing the surface expression levels of HLA-A, -B, and -C (Figures S4H and S4I). Collectively, these results confirm the feasibility of using the JI domain-containing reporters as effective tools for screening MHC class I-targeting viral antagonists.

### Screening of the MHC class I-targeting SARS-CoV-2 antagonist

Previous studies have demonstrated that both T cell numbers and T cell-mediated immunity are suboptimal in patients with severe COVID-19^19,20^. Our earlier work identified SARS-CoV-2 ORF6 as a major viral inhibitory factor targeting the MHC class I pathway^4^. To further explore additional SARS-CoV-2 viral inhibitors targeting the IRF1-NLRC5 axis-mediated MHC class I pathway, we utilized the JI-GFP reporter system (Figure 4A). Flow cytometry analysis revealed that SARS-CoV-2 viral proteins inhibited IRF1-induced JI-GFP reporter activity to varying degrees (Figures 4B and S5). Interestingly, several factors, including nonstructural protein 3 (Nsp3), ORF3a, ORF3d, ORF6, ORF7a, ORF7b, spike (S), and envelope (E) proteins, exhibited inhibitory effects exceeding threefold relative to the control (Figure 4B, red bar). These results were further validated using real-time quantitative PCR (RT-qPCR) to evaluate the expression levels of MHC class I-associated genes (Figure 4C).

To investigate the mechanistic basis of this phenotype, we examined whether these viral factors could suppress IRF1 nuclear translocation in response to IFNγ stimulation. Notably, two viral factors, ORF6 (as confirmed in our earlier study^4^) and ORF7b, significantly inhibited IRF1 nuclear localization (Figures 4D and 4E), identifying ORF7b as another SARS-CoV-2 antagonist targeting the IFNγ-IRF1 axis. Additionally, flow cytometry analysis confirmed that both ORF6 and ORF7b dramatically suppressed MHC class I surface expression (Figure 4F).

## DISCUSSION

NLRC5 and CIITA are the primary transcriptional regulators of MHC class I and MHC class II, respectively, and are essential for the adaptive immune response. Deficiency in either gene leads to severe clinical manifestations, including increased susceptibility to infections, autoimmune disorders, and cancer^21,22^. Therefore, the precise regulation of both genes is critical for maintaining immune homeostasis. While the regulatory mechanisms governing CIITA expression have been extensively characterized^14,15^, the fundamental transcriptional regulation of NLRC5 remains poorly understood. Although a previous study reported two *cis*-acting elements within the NLRC5 promoter region, we were unable to reproduce these findings in our system (Figure 2), suggesting the presence of yet unidentified regulatory elements.

Through a detailed dissection analysis of the NLRC5 promoter region, we identified two novel conserved *cis*-regulatory elements, an E-box and an ISRE, whose DNA sequences are completely conserved across various mammalian species. Notably, both elements play critical roles in NLRC5 transcriptional regulation: the E-box is required for basal promoter activity, whereas the ISRE is specifically involved in IFN-dependent upregulation of NLRC5 expression (Fig. 3).

Together with NLRC5, IRF1 plays a pivotal role in inducing the MHC class I pathway by directly binding to ISREs within MHC class I promoter regions^1^. Importantly, our study further elucidates that IRF1 contributes to MHC class I activation through two distinct mechanisms: functioning as a direct transcriptional regulator of MHC class I and enhancing CITA function by upregulating the NLRC5 expression at the transcriptional level.

IRF1 regulates the expression of various interferon-stimulated genes (ISGs) by recognizing specific DNA motifs^23^. One well-characterized ISG that IRF1 critically regulates is guanylate- binding protein (GBP). Particularly, IFNγ-mediated GBP induction is highly dependent on IRF1^24^. Interestingly, the consensus IRF1 binding sequence in the GBP promoter region, CTTTCAGTTTC, is identical to that of NLRC5 (Figure 3A). Based on this, it is plausible that NLRC5 expression is more strongly induced by IFNγ than by type I IFNs (IFNα/β). This could, in turn, explain why IFNγ has a more pronounced effect on the MHC class I pathway compared to type I IFNs^25,26^. However, this intriguing possibility remains to be further investigated.

Using the NLRC5 ISRE, we established an efficient system for screening agonists and antagonists that specifically regulate the IFNγ-IRF1-NLRC5 axis. Our screening tool successfully confirmed the inhibitory effects of previously reported viral antagonists (Figure 4). Moreover, through this system, we identified several novel SARS-CoV-2 antagonists that target the IFNγ- IRF1-NLRC5 pathway. Consistent with our previous study demonstrating that SARS-CoV-2 ORF6 inhibits the MHC class I pathway, our screening data further corroborate the inhibitory effect of ORF6. Additionally, we identified ORF7b as a novel viral factor that disrupts IFNγ-induced IRF1 nuclear translocation. While other SARS-CoV-2 proteins identified in this study were also found to suppress the IFNγ-IRF1-NLRC5 axis, their precise molecular mechanisms remain to be further elucidated.

Collectively, our study fills a critical gap in understanding the NLRC5-mediated antigen presentation pathway by elucidating the precise molecular mechanism of IFNγ-mediated NLRC5 expression. Furthermore, we demonstrate that JI-containing reporters serve as robust and reliable tools for screening antagonists targeting the IRF1-NLRC5-MHC class I axis. These insights and methodologies may facilitate the development of MHC class I-targeted therapeutics through precise modulation of the IRF1-NLRC5 axis.

## METHOD DETAILS

### Cells

HeLa, HEK293T, and VERO E6 cells were obtained from ATCC. IFNAR1 KO HeLa cells were a generous gift from Dr. Takashi Fujita (Kyoto University, Japan), and PH5CH8 WT and IRF1 KO cells were kindly provided by Dr. Yamane Daisuke (Tokyo Metropolitan Institute of Medical Science, Japan). All cells were maintained at 37 °C in a humidified incubator with 5% CO₂ in Dulbecco’s Modified Eagle Medium (DMEM) supplemented with 10% fetal bovine serum (FBS) and penicillin-streptomycin (100 U/ml and 100 µg/ml, respectively).

### RNA transfection

Poly I∶C (Cat. #tlrl-pic) was purchased from InvivoGen. 5’ ppp-RNA was synthesized by *in vitro* transcription using the EZ^TM^ High Yield In Vitro Transcription Kit (enzynomics, Cat. #EZ026S), as previously described^27^. 1 μg of RNA was delivered into the cells using Lipofectamine 2000, following the manufacturer’s instructions (Invitrogen, Cat. #11668019).

### Plasmids

The pGL3 plasmid was obtained from Promega. SARS-CoV-2 genes cloned into the pEF-BOS plasmid were kindly provided by Dr. Dae-Gyun Ahn (Korea Research Institute of Chemical Technology, Korea). Additionally, viral genes, including HCV Core and HBV X, were chemically synthesized and inserted into the pEF-BOS plasmid. The KSHV K11 expression plasmid was generously provided by Dr. Hye-Ra Lee (Korea University, Korea). The pEF-BOS-STAT1 vector was obtained from Dr. Takashi Fujita (Kyoto University, Japan), and the pEF-BOS-RIG-I CARD and pEF-BOS-MDA5-CARD plasmids were generous gifts from Dr. Mitsutoshi Yoneyama (Chiba University, Japan). Human IRF1, IRF3, and IRF7 were cloned into the pEF-BOS vector, and the IRF1 2WA and IRF1 lacking DNA binding domain mutant was generated using site-directed mutagenesis via PCR with specifically designed primer sets. Reporter constructs used in this study were generated by PCR amplification of DNA fragments from HeLa cell genomic DNA, using primer sets targeting the desired promoter regions. These fragments were subsequently inserted into either the pGL3 vector (for the luciferase-based system) or the pNifty-GFP vector (for the GFP-based system, kindly provided by Dr. Ella Sklan, Tel Aviv University, Israel).

### Flow cytometry

For the analysis of surface HLA expression, 3 × 10^5^ HEK293T cells were transfected with 1 μg of either empty or IRF1 WT constructs, along with the indicated viral antagonists. After 48 hours of stimulation, cells were washed with PBS, collected in FACS buffer (2% FBS/PBS), and stained with either a PE-conjugated isotype control (BioLegend, Cat. #400214) or a PE-conjugated anti-human HLA-A, B, C antibody (BioLegend, Cat. #311406). Following a 30-minute incubation on ice, cells were washed twice with FACS buffer and fixed with 4% paraformaldehyde (PFA) for 30 minutes at 4 °C. The fixed cells were then washed twice with FACS buffer and analyzed by flow cytometry using a FACS Canto (BD). Data analysis was conducted using FlowJo software (BD). To screen viral antagonists targeting the IRF1-NLRC5 axis, 3 × 10^5^ HEK293T cells were transfected with 1 μg of either an empty or JI-GFP construct, along with the indicated viral antagonists. After 48 hours of stimulation, cells were washed with PBS, collected in FACS buffer, and analyzed by flow cytometry as described above to measure GFP intensity.

### Electrophoretic mobility shift assay

For the analysis of NLRC5 ISRE and transcription factor interactions, a chemiluminescent nucleic acid detection kit (Thermo Scientific, Cat. #89880) was used according to the manufacturer’s protocol. Briefly, 0.6 μM of a biotin-labeled probe containing either the NLRC5 WT (5’- CTACTTTCAGTTTCGTGCAGAGCGC-3’) or mutant (5’- CTACGTGCAGGTGAGTGCAGAGCGC-3’) ISRE region was incubated with HEK293T cell lysates overexpressing IRF1, IRF3, or STAT in the reaction buffer provided by the manufacturer, supplemented with 50% glycerol, 100 mM MgCl2, and 1% NP-40. For the competition assay, a 15-fold excess of the unlabeled (cold) probe was added to the reaction. To assess ISRE-protein super-shift, 1 μg of the anti-IRF1 (Cell Signaling, Cat. #8478), anti-IRF3 (kindly provided by Dr. Takashi Fujita), and anti-STAT1 (Cell Signaling, Cat. #14994) antibodies were included in the reaction as indicated. After incubation at 25 °C for 30 min, the reaction mixtures were resolved by electrophoresis on a 7.5% non-denaturing polyacrylamide gel in 0.5× Tris-borate-EDTA (TBE) buffer at 4 °C. Following electrophoresis, immunoblotting was performed according to the manufacturer’s protocol, and the chemiluminescent signal was detected using ImageQuant™ LAS 500 (GE Healthcare).

### Immunofluorescence assay

HEK293T, HeLa WT, or IFNAR1 KO cells were plated on coverslips overnight and transfected with the indicated DNA constructs using Lipofectamine 2000 (Invitrogen, Cat. #11668019). For the JI-GFP reporter study, GFP intensity was monitored using fluorescent microscopy (Olympus) after 36 hours of transfection. For screening SARS-CoV-2 viral antagonists, cells were stimulated with IFNγ (100 U/ml, Peprotech, Cat. #300-02-100UG) for 7 hours after 36 hours of the indicated plasmid transfection. The cells were then fixed with 4% PFA for 20 minutes at 4°C, permeabilized with 0.05% Triton X-100 in PBS for 5 minutes at room temperature, blocked with 5 mg/ml BSA in PBST (0.04% Tween 20 in PBS) for 30 minutes, and incubated with the relevant primary antibodies diluted in blocking buffer at 4°C overnight. Following this, cells were incubated with secondary antibodies at room temperature for 1 hour. Nuclei were stained with Hoechst 33342 solution (Thermo, Cat. #62249) and analyzed using a confocal laser microscope (Olympus).

### Quantitative reverse-transcription PCR

For the evaluation of mRNA expression, total RNA was reverse transcribed into cDNA using the ReverTra Ace™ qPCR RT Master Mix reagent (Toyobo, Japan, Cat. #FSQ-301). RT-qPCR was then performed using THUNDERBIRD™ Next SYBR qPCR Mix reagent (Toyobo, Japan, Cat. #QPX-201) with specific primer sets targeting the indicated genes. The human GAPDH gene was used as an internal control for normalization. Relative gene expression levels were calculated using the ΔΔCt method, and results are presented as fold induction relative to untreated controls. The sequences of the specific primer sets used for RT-qPCR are provided in the STAR protocol.

### Reporter assay

For luciferase assays, 1 × 10^4^ cells were seeded into 48-well plates. To screen the cis-acting elements of the NLRC5 promoter, 50 ng of reporter constructs were co-transfected with 500 ng of either an empty plasmid or plasmids encoding human IRF1, RIG-I CARD, or MDA5 CARD genes. After 24 hours of incubation, cells were either transfected with 250 ng of 5’-ppp RNA or treated with 100 U/mL IFNβ (PeproTech, Cat. #300-02BC-5UG) or IFNγ (PeproTech, Cat. #300- 02-100UG) for an additional 12 hours. Cells were then harvested, and cell lysates were analyzed using the Luciferase Assay System (Promega).

### Prediction tools

To validate a CpG island within the NLRC5 promoter region (-4600 to +400 nucleotide position), MethPrimer (https://www.methprimer.com/cgi-bin/methprimer/methprimer.cgi) was utilized. For the analysis of the transcription start site of the NLRC5 gene, the DBTSS database (https://dbtss.hgc.jp/) was used for this study.

### Gene alignment

DNA sequences of NLRC5 promoter regions across various vertebrates: *Homo sapiens* (NG_030337), *Pan troglodytes* (XM_063797147), *Macaca mulatta* (XM_028840732), *Mus musculus* (AC128663), *Oryctolagus cuniculus* (XM_070061951), *Mustela putorius furo* (XM_045076089), *Bos taurus* (XR_011467716), *Capra hircus* (XR_011467716), *Ovis aries* (XM_060397955), *Equus caballus* (XM_023637031), *Camelus dromedarius* (XM_010996084), *Vicugna pacos* (XM_015248089), *Canis lupus familiaris* (XM_038659294), *Felis catus* (XM_019819771), *Cavia porcellus* (XR_010046976), *Desmodus rotundus* (XM_053916121), and *Tursiops truncatus* (XM_033845687), were aligned using CLUSTAL multiple sequence alignment via MUSCLE (3.8) software.

### Statistics

All statistical analyses were conducted using a two-tailed unpaired t-test. Statistical significance is denoted as *P < 0.05, **P < 0.01, ***P < 0.001, ****P < 0.0001, and ns indicates not significant. Data from three independent experiments were used. Error bars represent the mean ± SD. Graphs were generated using GraphPad Prism 10 software (GraphPad Software, USA).

## Acknowledgements

We thank Drs. Takashi Fujita (Kyoto University), Mitsutoshi Yoneyama (Chiba University), Dae- Gyun Ahn (Korea Research Institute of Chemical Technology, Korea), Hye-Ra Lee (Korea University), and Yamane Daisuke (Tokyo Metropolitan Institute of Medical Science, Japan) for reagents and cell lines. This work was supported by the National Research Foundation of Korea (NRF) grant funded by the Korea government (MSIT) (RS-2024-00407902) to J.-S. Y., Global - Learning & Academic research institution for Master’s·PhD students, and Postdocs (LAMP) Program of the National Research Foundation of Korea (NRF) grant funded by the Ministry of Education (No. RS-2023-00301914) to J.-S. Y., and Japan Society for the Promotion of Science (21K15445) to J.-S. Y.

## Author contributions

J.-S.Y. conducted the study, designed the experiments, analyzed the data, interpreted the results., and wrote the manuscript. J.-S.Y., Y.K, S.V, S.Y performed the experiments. K.S.K. contributed to data interpretation and discussion.

## Supplementary Figures

**Figure S1.**
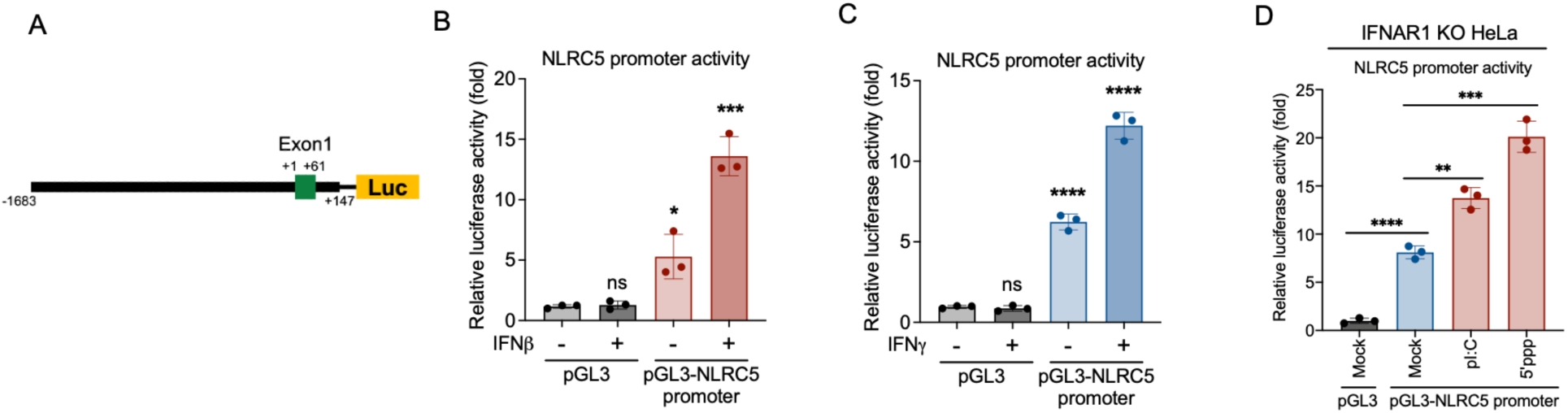
IFN-dependent and -independent NLRC5 promoter activation. (A) Schematic illustration of the luciferase-based NLRC5 promoter reporter construct, with the exact nucleotide position (-1683 to +147) indicated. (B and C) NLRC5 promoter activity was assessed by luciferase assay in HeLa cells treated with IFNβ (B) or IFNγ (C). (D) NLRC5 promoter activity was measured by luciferase assay in IFNAR1 KO HeLa cells under untreated conditions or stimulated with poly I:C (pI:C) or 5’-ppp RNA (5’ppp).

**Figure S2.**
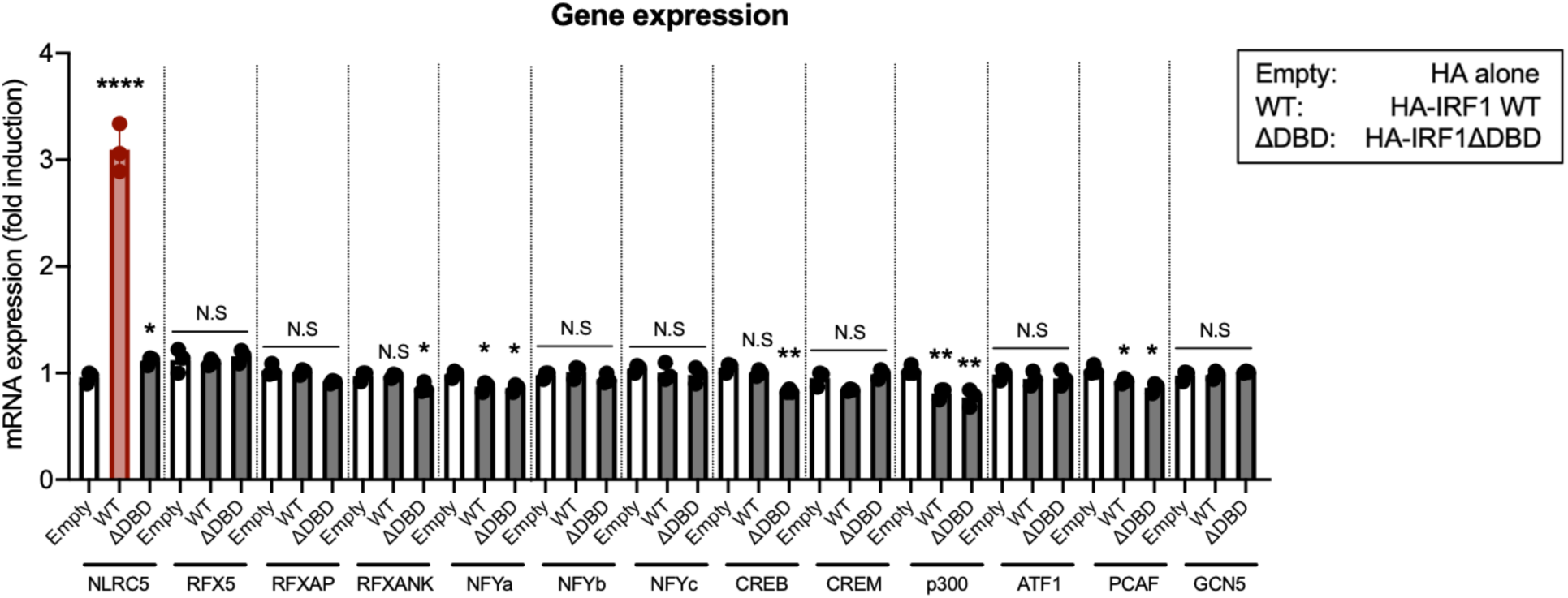
IRF1 specifically induces NLRC5 transcription The expression levels of all known transcription factors and transactivators involved in the induction of the MHC class I pathway were evaluated by RT-qPCR in HEK293T cells transfected with either an empty vector, IRF1 WT, or IRF1 lacking the DNA binding domain.

**Figure S3.**
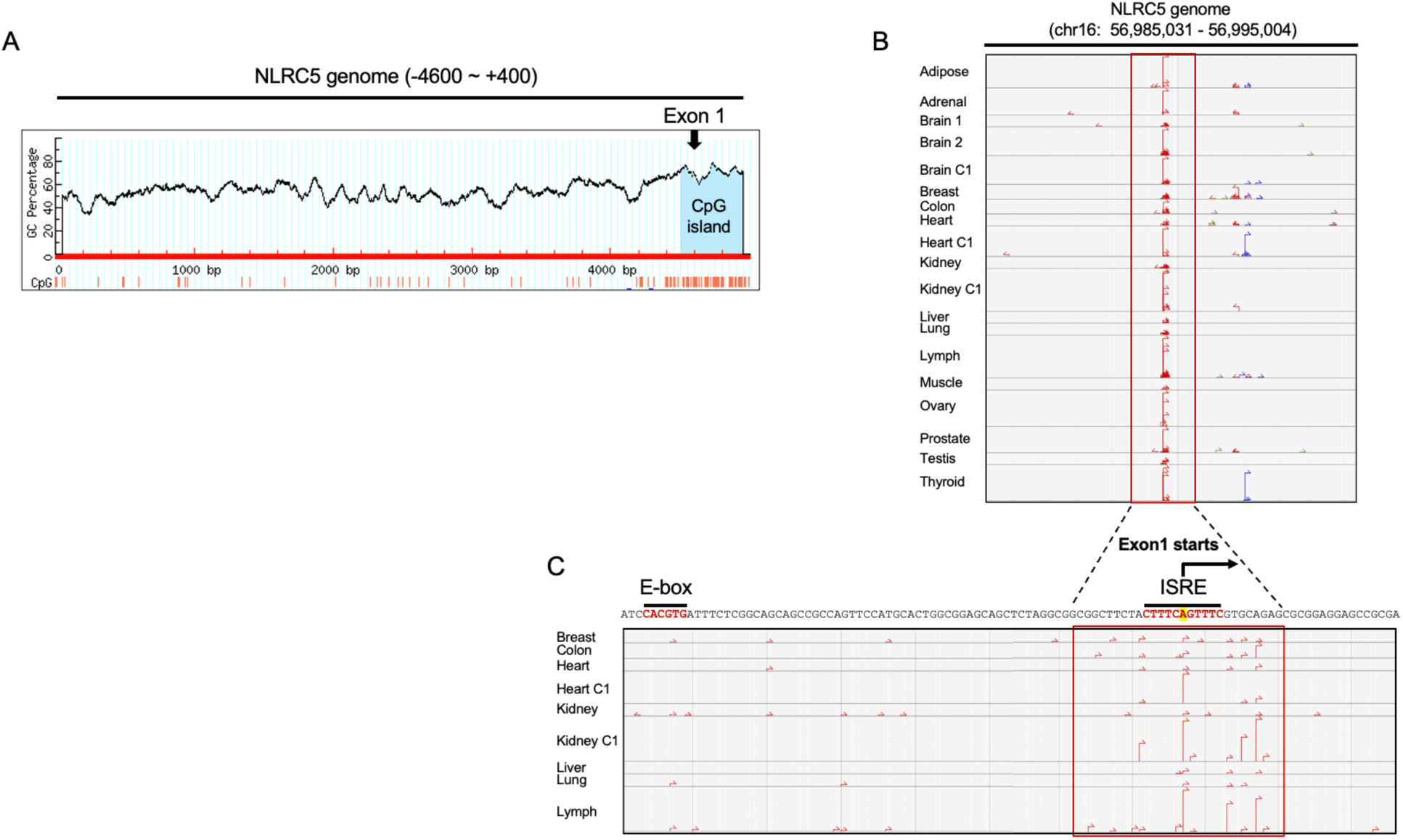
NLRC5 JI domain possesses a putative transcription starting site (A) The CpG island within the NLRC5 genomic region (-4600 to +400) was analyzed using the MethPrimer tool. The position of Exon 1 is indicated, and the CpG island is highlighted in blue. (B and C) Analysis of the NLRC5 transcription start site. Transcription start sites of NLRC5 across various tissues are indicated by red arrows (B), with an enlarged view provided in (C). The genomic coordinates of NLRC5 and the corresponding DNA sequences are shown. DNA sequences corresponding to the E-box and ISRE motifs are highlighted in red.

**Figure S4.**
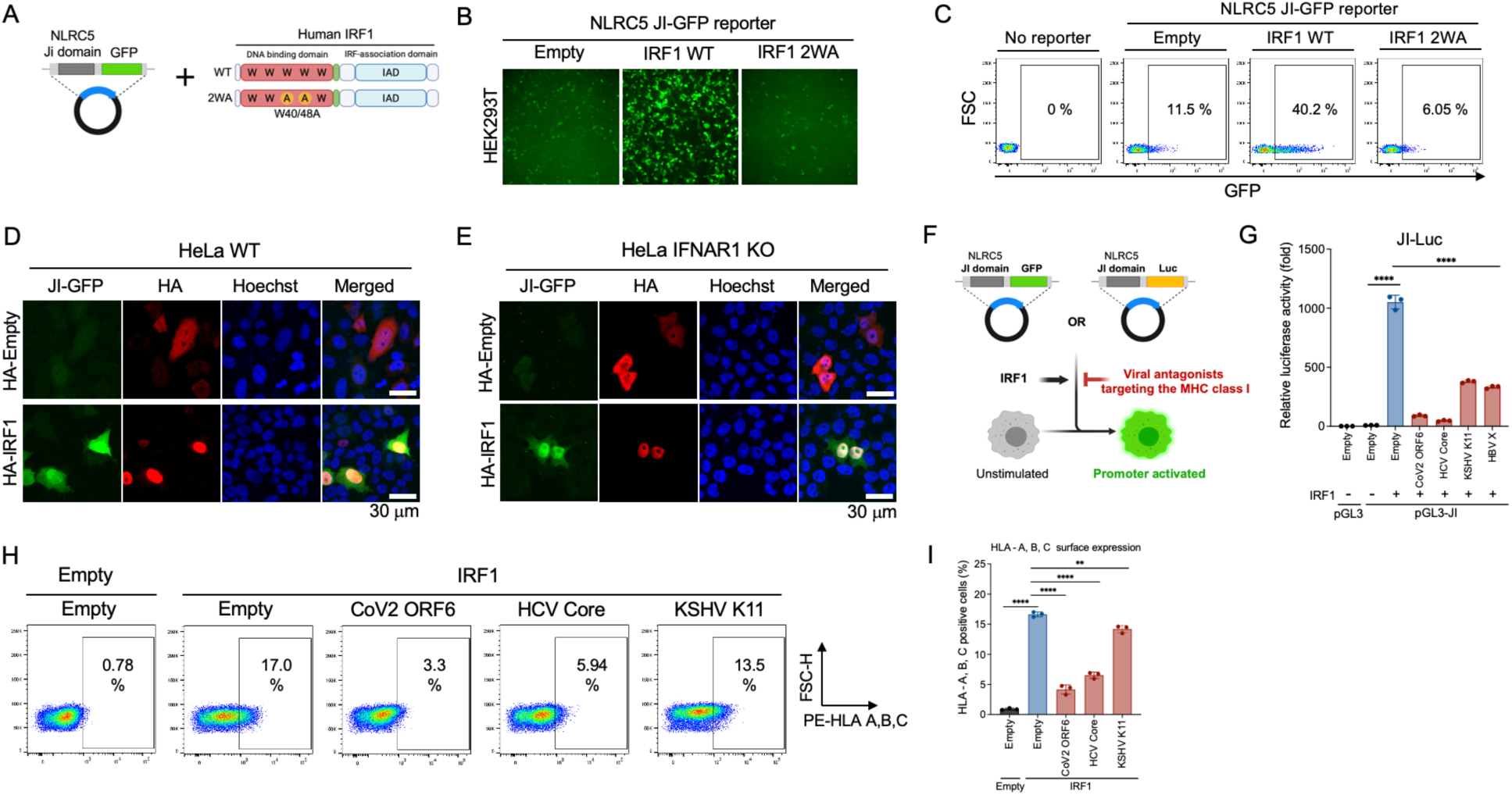
Establishment of GFP- and luciferased-based monitoring system for NLRC5 expression via NLRC5 JI domain (A) Schematic representation of the IRF1-mediated NLRC5 JI-GFP reporter system. (B and C) IRF1-driven NLRC5 JI-GFP reporter activity was assessed using fluorescence microscopy (B) and flow cytometry (C). (D and E) Activation of the NLRC5 JI-GFP reporter by IRF1 was analyzed via confocal microscopy in WT HeLa cells (D) and IFNAR1 KO HeLa cells (E). (F) Schematic representation of the strategy for validating viral antagonists targeting the IRF1-NLRC5 axis using the NLRC5 JI-GFP system. (G) The inhibitory effects of viral antagonists on IRF1-mediated NLRC5 JI-Luc activation were assessed using a luciferase assay. (H and I) The impact of viral factors on IRF1-induced surface expression of HLA-A, -B, and -C was evaluated by flow cytometry (H), with quantitative data presented as a bar graph (I).

**Figure S5.**
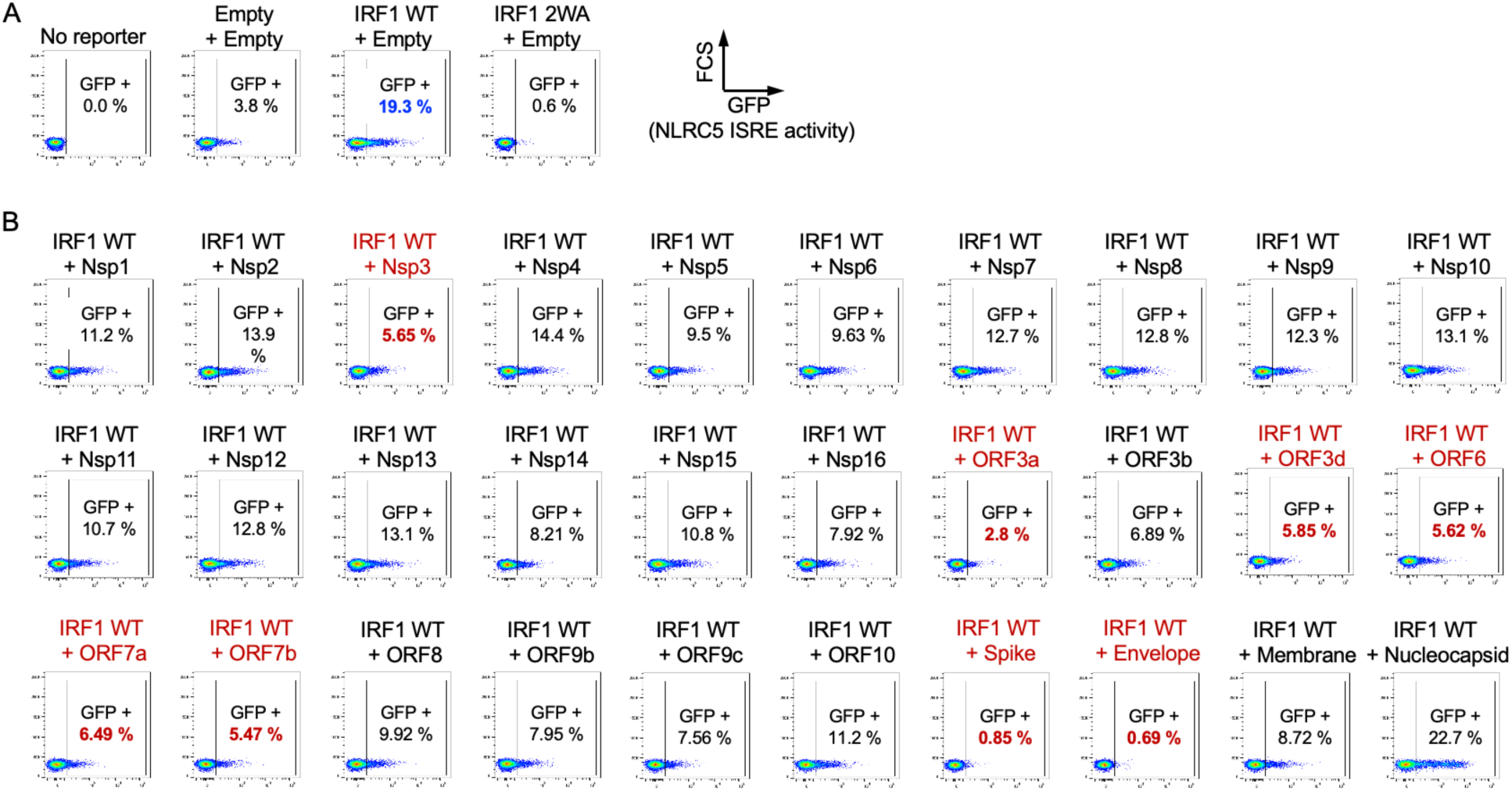
Identification of the SARS-CoV-2 factors antagonizing the IRF1-NLRC5 axis- derived MHC class I pathway (B) IRF1-mediated NLRC5 JI-GFP activation was validated using flow cytometry, with an empty vector and IRF1 2WA used as controls. (B) The inhibition of IRF1-driven NLRC5 JI-GFP reporter activation by SARS-CoV-2 viral proteins was assessed via flow cytometry. SARS-CoV-2 factors that led to more than a threefold reduction in activation are highlighted in red.

